# Confidence in global motion discrimination is better predicted by perceptual reliability than by stimulus-based heuristics

**DOI:** 10.64898/2026.11.12.766172

**Authors:** Can Oluk, Martin Szinte, Guillaume S. Masson, Pascal Mamassian

## Abstract

Decision-making requires evaluating the reliability of perceptual estimates, but whether confidence reflects perceptual reliability or stimulus-based heuristics remains debated. We used a multiple-aperture global motion stimulus in which the global direction, and therefore motion coherence, is only available after integrating locally ambiguous signals, so that no simple stimulus statistic can serve as a shortcut. Observers reported the global direction in two successive intervals and then chose which decision was more likely to be correct. Confidence tracked accuracy beyond what coherence alone could explain: observers were more accurate in chosen intervals even at the same coherence level. Observers also frequently reported the direction opposite to the true motion, and, in direct comparisons, were more confident in these errors than in other large errors. We compared two image-computable models of motion integration. A standard motion-energy model failed to produce opposite-direction errors and the associated confidence pattern. A normalized motion-energy model, adding tuned inhibition among direction channels and an untuned gain control that increases overall activation with coherence, accounted for most of the variance in both perceptual decisions and confidence judgments. Tuned inhibition, which must be sharp if it is close to excitation in strength, produced the opposite-direction errors and reproduced the direction of their confidence advantage, and the untuned gain was needed for confidence to depend on coherence as strongly as observed. Confidence was read out from the same signal as the perceptual decision, and noise at the confidence stage, although probably present, added little to the fit beyond explaining the accuracy difference between chosen and unchosen intervals. These results suggest that confidence in this task relies on perceptual reliability. They also suggest that seemingly unusual metacognitive behavior can emerge from perceptual computations, which must therefore be modeled explicitly to identify genuine confidence mechanisms.

**Author summary:** When we make a decision, we usually also know how likely it is to be right. This sense of confidence could come from evaluating how reliable our perception is, or from simple shortcuts, such as how clear or noisy the image looks. In most previous studies, these two sources were hard to separate. We used displays of hundreds of drifting patterns whose individual motions are ambiguous and only reveal a common direction once the brain combines them. Observers judged the direction of motion, and then which of two decisions was more likely to be correct. Their confidence tracked their accuracy beyond what motion strength alone could explain. Surprisingly, observers often reported the opposite direction and were more confident in these errors than in other large errors. A motion-processing model explained both with two mechanisms: neurons tuned to similar directions suppress each other, which produces the opposite-direction errors, and overall activity rises with the proportion of elements moving together, which makes confidence depend strongly on motion strength. These results suggest that confidence in this task relies on how reliable our perception is. They also suggest that seemingly strange confidence behavior can arise from how the visual system processes motion.

## Introduction

Human decision-making depends not only on accurately processing sensory information, but also on evaluating the reliability of the resulting perceptual estimates. Avoiding a predator, for example, requires estimating the speed and direction of its movement, but also judging how likely that estimate is to be correct in order to select the most appropriate response. Such confidence judgments play a key role in many cognitive functions, including guiding information seeking [1], controlling evidence accumulation [2], and enabling collective decision-making [3]. Accordingly, humans are generally able to report confidence that tracks the accuracy of their decisions across a wide range of conditions [4–6].

When forming these confidence judgments, observers can draw on different sources of information. A common assumption is that confidence reflects perceptual reliability, that is, the precision of the internal estimate on which the perceptual decision is based [7,8]. Alternatively, observers might rely on stimulus-based heuristics, using a readily accessible stimulus statistic as a direct proxy for confidence. In visual motion, for example, observers might base their confidence on the contrast or coherence of the motion signals.

The extent to which humans rely on perceptual reliability rather than on stimulus-based heuristics remains debated. Some studies have shown that observers can make accurate confidence judgments even when performance depends on multiple visual attributes [9], suggesting that confidence extends beyond what any single stimulus-based heuristic could provide. Other studies, however, have shown that confidence can be manipulated through stimulus statistics without affecting perceptual accuracy [10–12], suggesting that, at least in some contexts, confidence may be driven largely by factors other than perceptual reliability [13]. One explanation for these discrepancies is that most previous studies used simple stimuli in which a single statistic, such as contrast or noise, is a readily available proxy for reliability. Because such statistics usually covary with perceptual reliability, the two accounts make similar predictions. It is therefore important to consider more complex paradigms in which no low-level statistic can serve as a shortcut, enabling a precise assessment of the perceptual reliability hypothesis. Moreover, because stimulus statistics and perceptual reliability both arise from the same sensory processing, formally separating the two hypotheses requires computational models that tease apart how low-level stimulus statistics and mid-level perceptual computations each contribute to confidence formation.

Global motion integration meets both requirements. We investigated the computational mechanisms of human confidence judgments using multiple-aperture global motion stimuli [14]. The stimuli consist of 468 drifting Gabors, all identical in size, contrast and spatial frequency but with random orientations (Fig 1a). Unlike in random-dot kinematograms (RDKs), the motion direction of each element is ambiguous because of the aperture problem [15,16]: when viewed in isolation, each Gabor appears to move in the direction orthogonal to its orientation. When the elements are presented together and are consistent with a single global direction, however, the visual system combines these local signals into a coherent global motion percept, a process called 2D motion integration (Fig 1b). For such 1D Gabor arrays, a linear vector average of the local motion signals predicts the correct global direction but a strongly underestimated speed (about half the true speed when orientations are uniformly distributed), whereas the non-linear intersection-of-constraints (IOC) rule recovers both the veridical direction and speed. Based on measurements of perceived global speed, Amano et al. [14] showed that the perception of such stimuli follows IOC.

**Fig 1.**
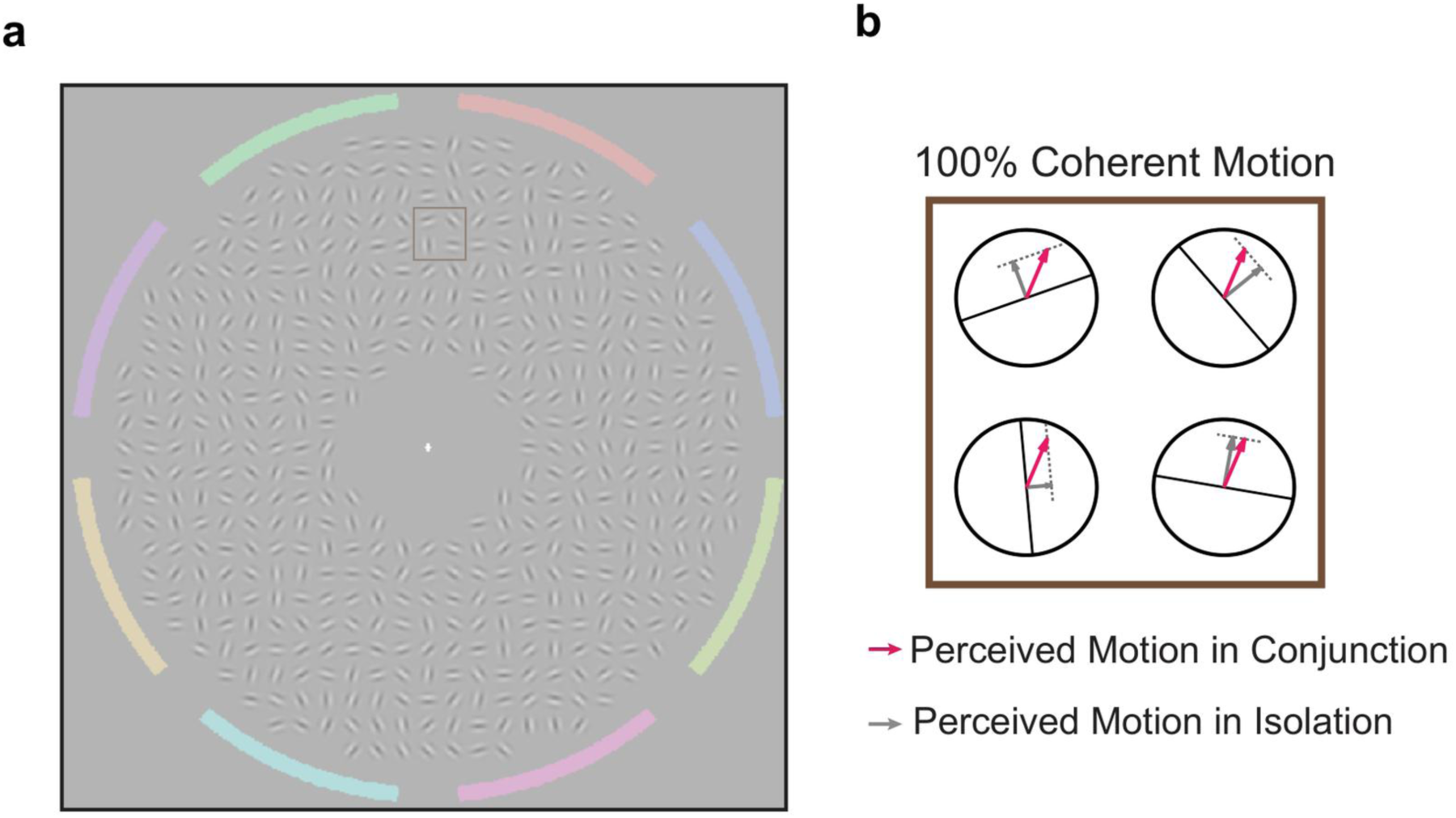
Multiple-aperture global motion stimulus. (a) A single frame of an example stimulus, consisting of 468 drifting Gabors with identical size, contrast and spatial frequency but random orientations. On each presentation, a proportion of elements (the motion coherence) drifts consistently with a single global 2D motion, whose speed is fixed and whose direction is one of eight possibilities. The remaining elements drift consistently with random global directions. The colored arcs of the surrounding ring indicate the eight response alternatives of the 8-AFC task. (b) Illustration of 2D motion integration for a subset of elements (square in a) at 100% coherence. Because of the aperture problem, the local motion of each element is ambiguous: viewed in isolation, a Gabor appears to move orthogonally to its orientation (grey arrows). All 2D velocities consistent with this local 1D motion lie on a constraint line parallel to the Gabor’s orientation (dashed lines [17]). When the elements are viewed together, the visual system recovers the single global motion consistent with all local signals (red arrows), which lies at the intersection of their constraint lines (intersection of constraints, IOC).

This stimulus design has two important consequences for confidence. First, because the global direction, and therefore motion coherence, is only available after the integration stage, no low-level stimulus statistic can serve as a shortcut to confidence. Previous work with similar stimuli has shown that confidence can be biased toward the clarity of local elements rather than global coherence [18]. In our stimuli, all elements had the same contrast and no added local noise, so local signal clarity provided no cue to confidence. Second, the design allows clear, competing predictions. If confidence is based on the integrated 2D motion signal, it should track perceptual performance, indicating access to perceptual reliability. If observers instead relied on unintegrated local motion signals, confidence should be systematically mismatched with performance. The remaining candidate heuristic, using the integrated motion coherence itself as a proxy for confidence, makes its own distinct predictions, which we test directly.

In the present study, we manipulated motion coherence by varying the proportion of Gabors whose local motion was consistent with a single global direction. On each trial, participants reported the global motion direction in two successive intervals (eight-alternative forced choice) and then indicated which of their two decisions was more likely to be correct (Fig 2; confidence forced choice [19]). To relate these judgments to the underlying computations, we modeled both decisions with motion-energy models of 2D motion integration [20,21], which account for a range of behavioral [22] and neurophysiological [23] findings. Because running such models at the pixel level for hundreds of elements is computationally prohibitive, we adapted the efficient image-computable approximation developed by Kane et al. [24]. We compared a standard motion-energy (SME) model, in which both the perceptual decision and confidence are read out directly from template-matched motion energy, with a normalized motion-energy (NME) model that adds two biologically motivated mechanisms: tuned inhibition among direction channels (motivated by the strong suppressive inputs to MT neurons [23,25]) and an untuned gain control that scales overall activation with motion coherence [26,27]. In both models, confidence is computed from the same signal that drives the perceptual decision, without a separate heuristic. As an additional analysis, we tested whether noise at the confidence stage improves the fit. We found that the NME model fitted the data better than the SME model and, with only these two perceptually inspired mechanisms, accounted for most of the variance in both perceptual decisions and confidence judgments, including the frequent opposite-direction errors and their relatively high confidence in direct comparisons, suggesting that confidence reflects the reliability of the integrated motion signal.

**Fig 2.**
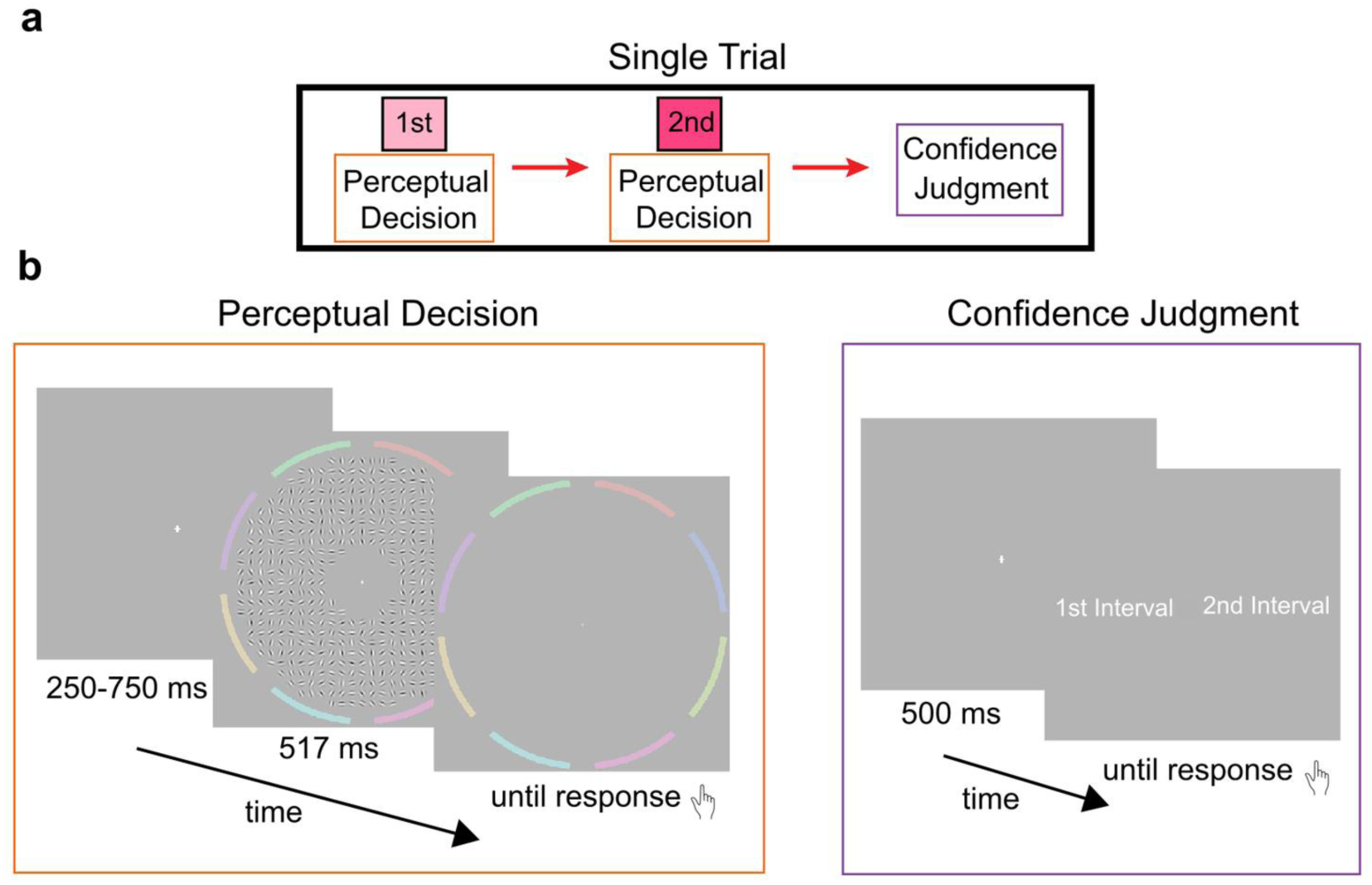
Experimental design. (a) A single trial comprises two perceptual decision intervals followed by a confidence judgment. In each perceptual interval, participants view a motion stimulus with independently sampled motion coherence and global direction, and report the perceived global motion direction. (b) Left: each perceptual decision begins with a jittered fixation period (250–750 ms), followed by a 517 ms motion stimulus enclosed by the eight-segment colored response ring. After motion offset, the ring remains visible until the participant selects the perceived direction with a mouse click. Right: for the confidence judgment, a 500 ms fixation period is followed by a prompt. Participants indicate which interval’s decision they believe was more likely to be correct by pressing the corresponding arrow key (left for the first interval, right for the second).

## Results

### Experimental Results

#### Perceptual decisions

We first evaluated direction discrimination accuracy and the distribution of perceptual errors. Averaged across coherence levels, the average observer identified the correct global motion direction on 65% of trials, with considerable variability across individuals (50–81%; chance level 12.5%). As expected, accuracy increased with motion coherence (Fig 3a): a bootstrapped linear fit gave a slope of 0.86 per unit coherence (95% CI [0.80, 0.93]).

**Fig 3.**
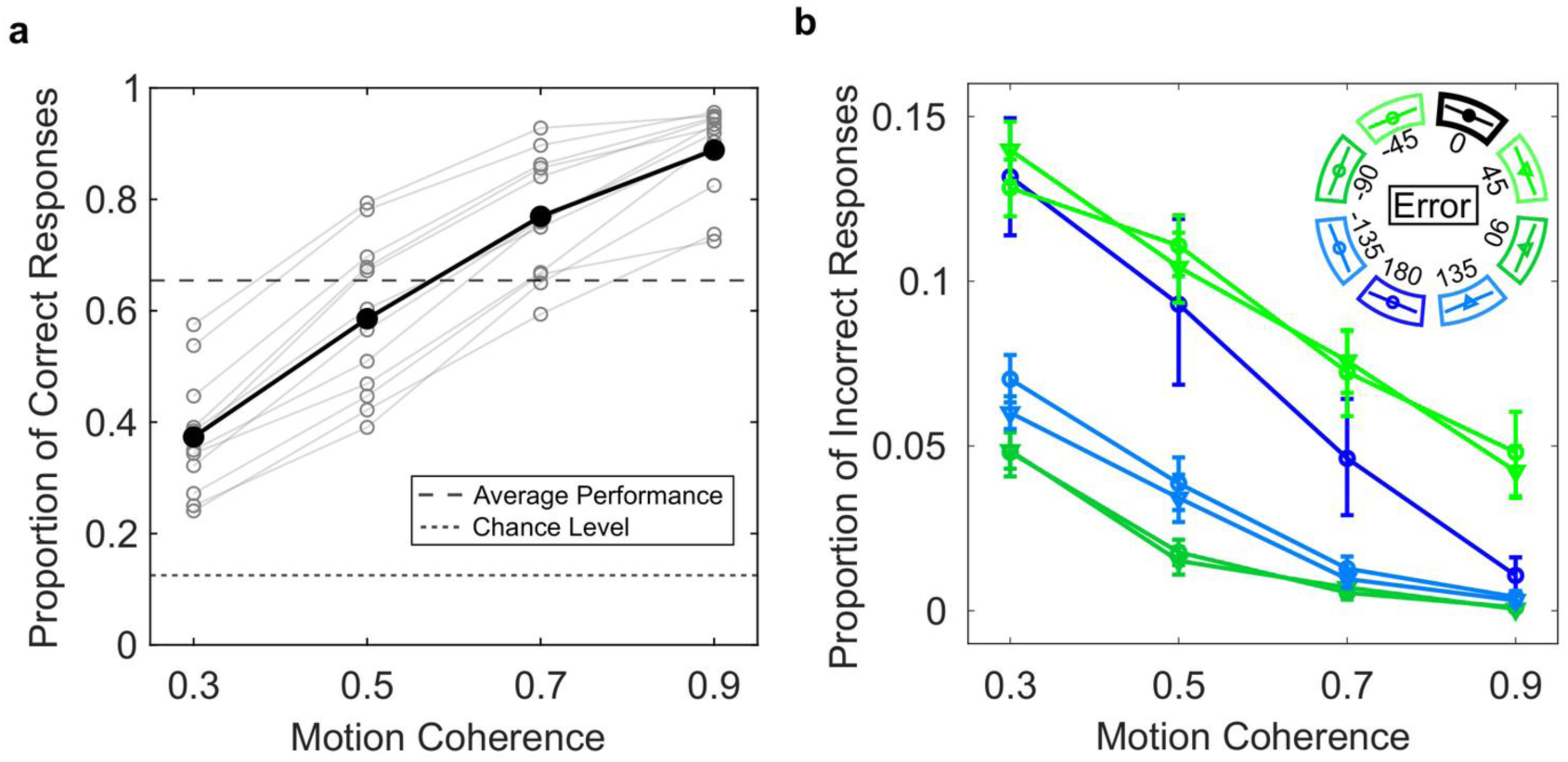
Perceptual decisions. (a) Proportion of correct responses as a function of motion coherence. Gray lines show individual participants and the black line shows the average observer. The dotted line indicates chance level (0.125), and the dashed line indicates the overall proportion correct of the average observer (0.65). (b) Proportion of incorrect responses as a function of motion coherence, for each response direction relative to the true motion direction (0°; see inset). Proportions are shown per response alternative: ±45° (light green), ±90° (dark green), ±135° (light blue) and 180° (dark blue). The two alternatives at the same angular distance (e.g., +45° and −45°) are plotted as separate lines of the same color. At each coherence level, the proportions of correct and incorrect responses sum to one. Error bars show 68% bootstrapped confidence intervals across participants.

We next examined the distribution of incorrect responses (Fig 3b). Responses were averaged across the eight true directions and expressed relative to the true direction (0°), so that, at each coherence level, the proportions of correct and incorrect responses sum to one. Because there are two alternatives at ±45°, ±90° and ±135° but only one at 180°, we compare error rates per response alternative. As expected, the most common error was to choose one of the two directions adjacent to the true direction (±45° errors, light green). Surprisingly, the next most common error was to choose the direction opposite to the true motion (180° errors, dark blue). Bootstrapped comparisons confirmed that 180° errors were more frequent than ±135° errors (light blue; mean difference = 0.04, 95% CI [0.02, 0.07]) and ±90° errors (dark green; mean difference = 0.05, 95% CI [0.03, 0.09]). The least frequent errors were those perpendicular to the true direction (±90°), which occurred less often than ±135° errors (mean difference = 0.011, 95% CI [0.006, 0.016]).

All error rates decreased as motion coherence increased, but 180° errors decreased more steeply than the other large errors. Pairwise comparisons of bootstrapped slopes confirmed that the decline in 180° errors was steeper than that of ±135° errors (Δslope = −0.10, 95% CI [−0.14, −0.06]) and ±90° errors (Δslope = −0.13, 95% CI [−0.18, −0.08]). As a consequence, at low coherence, opposite-direction errors were nearly as frequent as errors to the adjacent directions, whereas at high coherence they became almost as rare as ±90° and ±135° errors. This pattern was consistent across participants (S1 Fig).

#### Confidence judgments

For the confidence judgment, participants chose which of their two perceptual decisions was more likely to be correct. For the average observer, first- and second-interval choices were nearly balanced (47% vs. 53%). Only three participants deviated from an even split by more than 10 percentage points (first-interval choices: 37%, 73% and 13%). Such idiosyncratic interval biases have been reported previously and are stable within participants [11]; we account for them in the models with an interval-bias parameter.

To estimate metacognitive sensitivity, we classified each perceptual decision as selected (chosen as more likely to be correct) or unselected, based on the confidence judgment of that trial. Accuracy was 75% (95% CI [69%, 81%]) in selected intervals and 56% (95% CI [50%, 62%]) in unselected intervals, above and below the overall accuracy of 65%, respectively. This advantage of 19 percentage points (95% CI [16, 22]) was present in every participant, indicating that observers could monitor their own perceptual performance. Because selected intervals also tended to have higher motion coherence, part of this advantage could arise from coherence alone; we address this possibility below.

Building on this basic sensitivity, we next examined how confidence judgments relate to the magnitude of perceptual errors. We grouped errors into four categories: no error (0°), small error (±45°), large error (±90° and ±135°) and opposite error (180°). Depending on the combination of errors in the two intervals, each trial fell into one of 16 cases (4 × 4). Fig 4a shows the proportion of first-interval confidence judgments for each case: the x-axis gives the error category in the first interval, and line colors give the error category in the second interval. Marker size is proportional to the number of trials per case; 44% of trials contained no error in either interval (large black marker on the left). S2 Fig shows the same data in a different format, as matrices of choice proportions for every combination of error categories and of coherence levels in the two intervals, together with the predictions of the two models described below.

**Fig 4.**
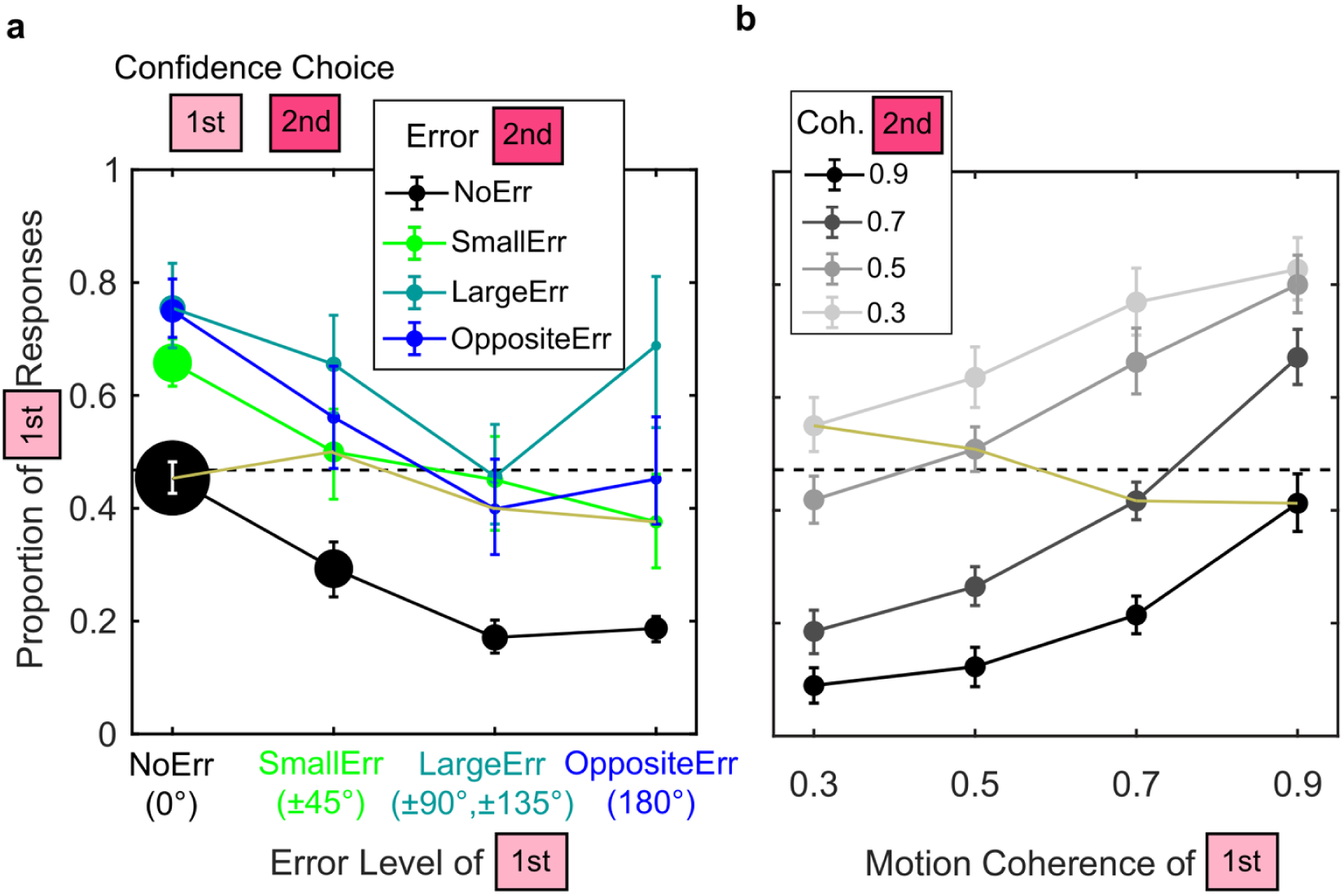
Confidence judgments. (a) Proportion of first-interval confidence judgments as a function of the perceptual error in the first interval. Line colors indicate the error in the second interval: no error (0°, black), small error (±45°, green), large error (±90° and ±135°, cyan) and opposite error (180°, blue). Marker size is proportional to the number of trials in each case; error bars on the largest marker are drawn in white for visibility. (b) Proportion of first-interval confidence judgments as a function of motion coherence in the first interval. Shades of gray indicate motion coherence in the second interval, from black (0.9) to light gray (0.3). In both panels, the yellow line connects conditions in which the two intervals are equal (same error category in a, same coherence in b), the dashed horizontal line marks the overall proportion of first-interval choices (0.47), and error bars show 68% bootstrapped confidence intervals across participants.

When the two intervals contained errors of the same category (yellow line in Fig 4a), the proportion of first-interval choices stayed close to the overall average (dashed line). Overall, confidence judgments tracked perceptual errors: the larger the error in the first interval, the less often observers chose it (lines decrease from left to right), and the larger the error in the second interval, the more often they chose the first interval (lines shift upward from black to green to cyan). Opposite-direction errors were a striking exception to this pattern. When the second interval contained a 180° error (blue line), observers chose the first interval less often than when the second interval contained a large error (cyan line). Likewise, along the cyan line, observers chose the first interval more often when it contained a 180° error than when it contained a large error. When an opposite error was paired with a large error, observers chose the opposite-error interval in 60% (68% CI [50%, 68%]) of trials when it was the second interval and in 69% (68% CI [55%, 82%]) when it was the first. Pooled across the two orders (119 trials), the opposite-error interval was chosen in 64% of trials (68% CI [57%, 71%]). Of the 11 participants with such trials, 9 chose the opposite-error interval more often than the large-error interval, one chose them equally often and one less often (S3 Fig). Thus, opposite-direction errors were not only frequent but also, when directly compared, associated with higher confidence than other large errors (±90° and ±135°).

Next, we asked whether confidence judgments reflect perceptual reliability or a stimulus-based heuristic relying on motion coherence. The proportion of first-interval choices increased systematically with the difference in coherence between the two intervals (Fig 4b), and this relationship was consistent across participants (S4 Fig). This pattern is predicted by both accounts: lower coherence reduces perceptual reliability, but observers could also use the coherence of the stimulus directly as a proxy for confidence.

To distinguish these accounts, we simulated confidence judgments driven entirely by coherence: in each trial, the interval with the higher coherence was selected, and accuracy depended only on coherence. This heuristic reproduced the overall accuracy difference between selected and unselected intervals (76% vs. 55% in the simulation, compared to 75% vs. 56% in the data). However, it makes a strict prediction: at any given coherence level, selected and unselected intervals must have the same accuracy, so that their curves overlap (Fig 5b). The data violate this prediction (Fig 5a). Averaged across coherence levels, observers were more accurate in selected than in unselected intervals (mean difference = 7.8 percentage points, 95% CI [6.2, 9.5]). Confidence therefore did not depend on the nominal coherence level alone, but tracked trial-to-trial fluctuations in the strength of the integrated motion signal, that is, perceptual reliability. Because coherence in our stimuli can only be estimated after motion integration, even a heuristic based on perceived coherence would require access to this integrated signal. We formalize this account with the computational models below.

**Fig 5.**
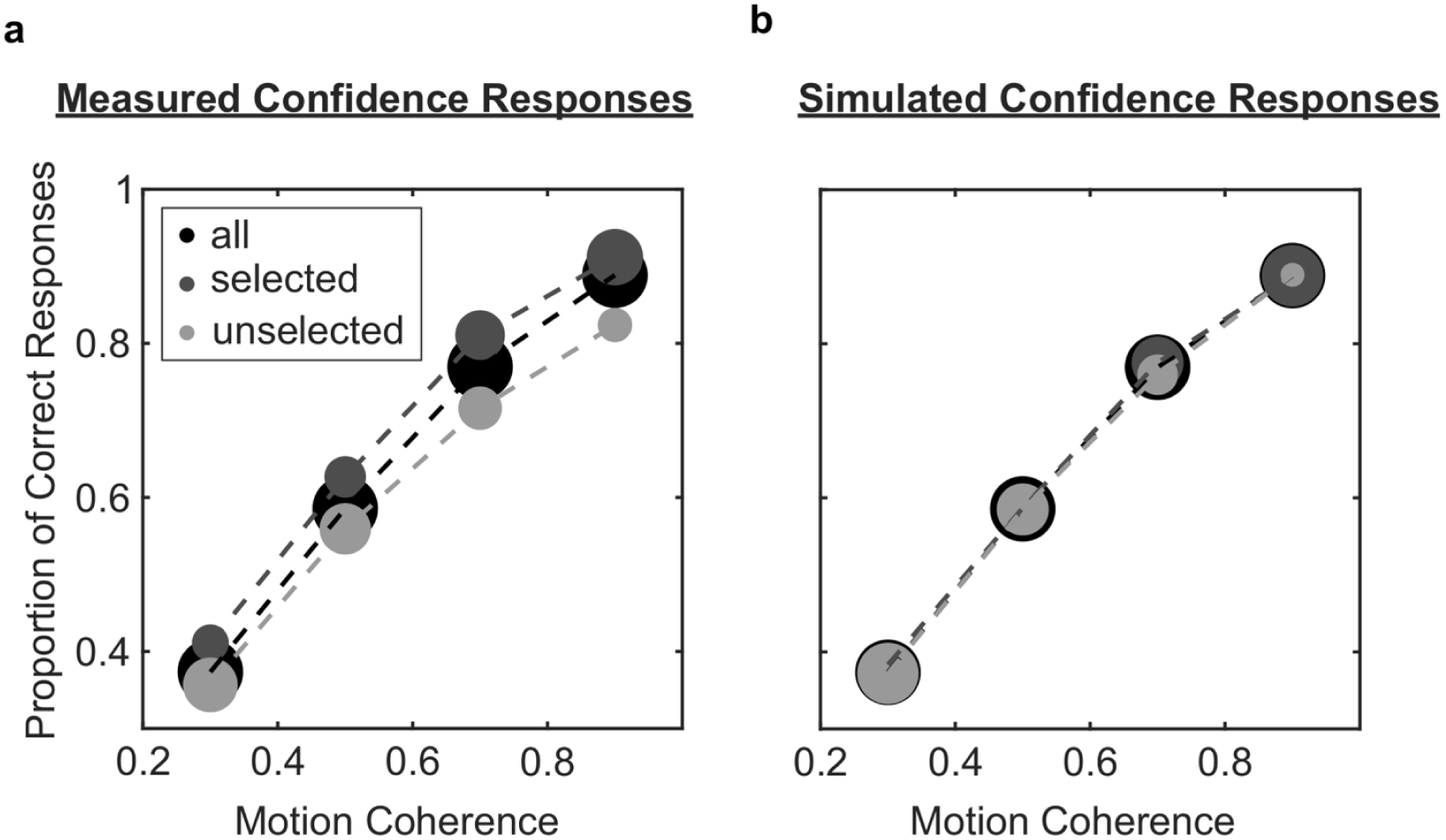
Perceptual accuracy separated by confidence judgment. Proportion of correct responses as a function of motion coherence for all intervals (thin dashed black line), selected intervals (dark gray; chosen as more likely to be correct) and unselected intervals (light gray). Marker size is proportional to the average number of trials per condition. (a) Measured data. At every coherence level, observers were more accurate in selected than in unselected intervals. Error bars show 68% bootstrapped confidence intervals across participants. (b) Simulation of a coherence heuristic, in which the interval with the higher motion coherence is always selected (ties are resolved at random) and accuracy depends only on coherence. For each participant, simulated responses were drawn from that participant’s accuracy at each coherence level. Under this heuristic, the number of selected and unselected trials varies with coherence, but at any given coherence level, selected and unselected intervals have the same accuracy, so the curves overlap (up to simulation noise). The separation observed in (a) therefore cannot be explained by coherence alone.

Finally, we observed an order bias when the two intervals had the same coherence (yellow line in Fig 4b): observers chose the first interval more often at low coherence (30% and 50%) and the second interval more often at high coherence (70% and 90%). This bias was not accompanied by a corresponding difference in accuracy between intervals (see Data analysis), and is consistent with a contraction of the first-interval signal toward the average in memory, which we model explicitly below.

### Modeling Results

Our second goal was to develop a computational model that predicts both perceptual decisions and confidence judgments. For each trial, the model must estimate the perceived global motion direction and, at the same time, provide a measure of perceptual reliability. To this end, we adapted established models of direction-selective neurons tuned to global motion, often referred to as pattern cells [21,23]. We compared two models: a standard motion-energy (SME) model, based on spatial pooling of local motion energy and IOC templates (Fig 6), and a normalized motion-energy (NME) model, which adds tuned inhibition among direction channels and an untuned gain control that scales total activation with motion coherence (Fig 7).

**Fig 6.**
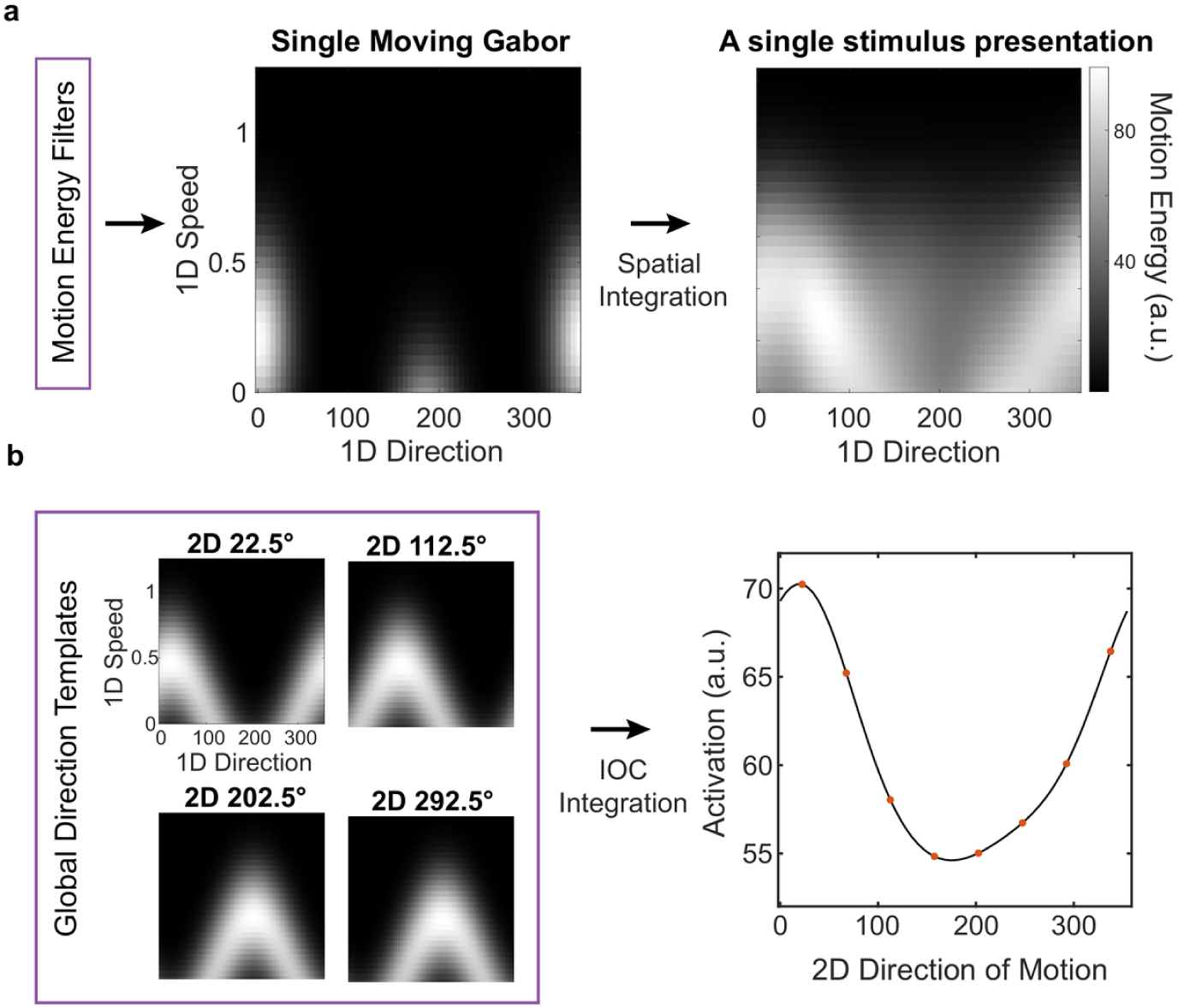
The standard motion-energy (SME) model. (a) Each moving Gabor is represented by its motion energy across 1D directions and speeds, approximated by a 2D Gaussian. Left: simulated motion energy of a single Gabor drifting at 0.21 dva/s in the 1D direction of 0°. The direction axis is circular, and energy at negative speeds wraps onto the opposite direction (180° away) at the corresponding positive speed. Right: total motion energy of a single stimulus presentation, obtained by summing the motion energy of all 468 Gabor patches across space. In this example, the stimulus has 30% coherence, a global direction of 22.5° and a global speed of 0.5 dva/s. (b) Global direction templates, corresponding to the motion-energy pattern of 100% coherent motion in each candidate direction (four of the eight templates are shown). Each template concentrates its energy along the half-cosine curve of 1D velocities consistent with one 2D motion, so that template matching implements the intersection of constraints (IOC) at the stimulus speed. The 22.5° template was obtained by averaging the motion-energy profiles of 700 presentations of 100% coherent stimuli moving at 0.5 dva/s; templates for the other directions were obtained by circularly shifting it along the direction axis, and all templates were normalized to sum to one. The dot product of each template with the total motion energy of the stimulus (a, right) gives the activation for that direction (right panel). The direction with the maximum activation is the perceived global direction (here, 22.5°).

**Fig 7.**
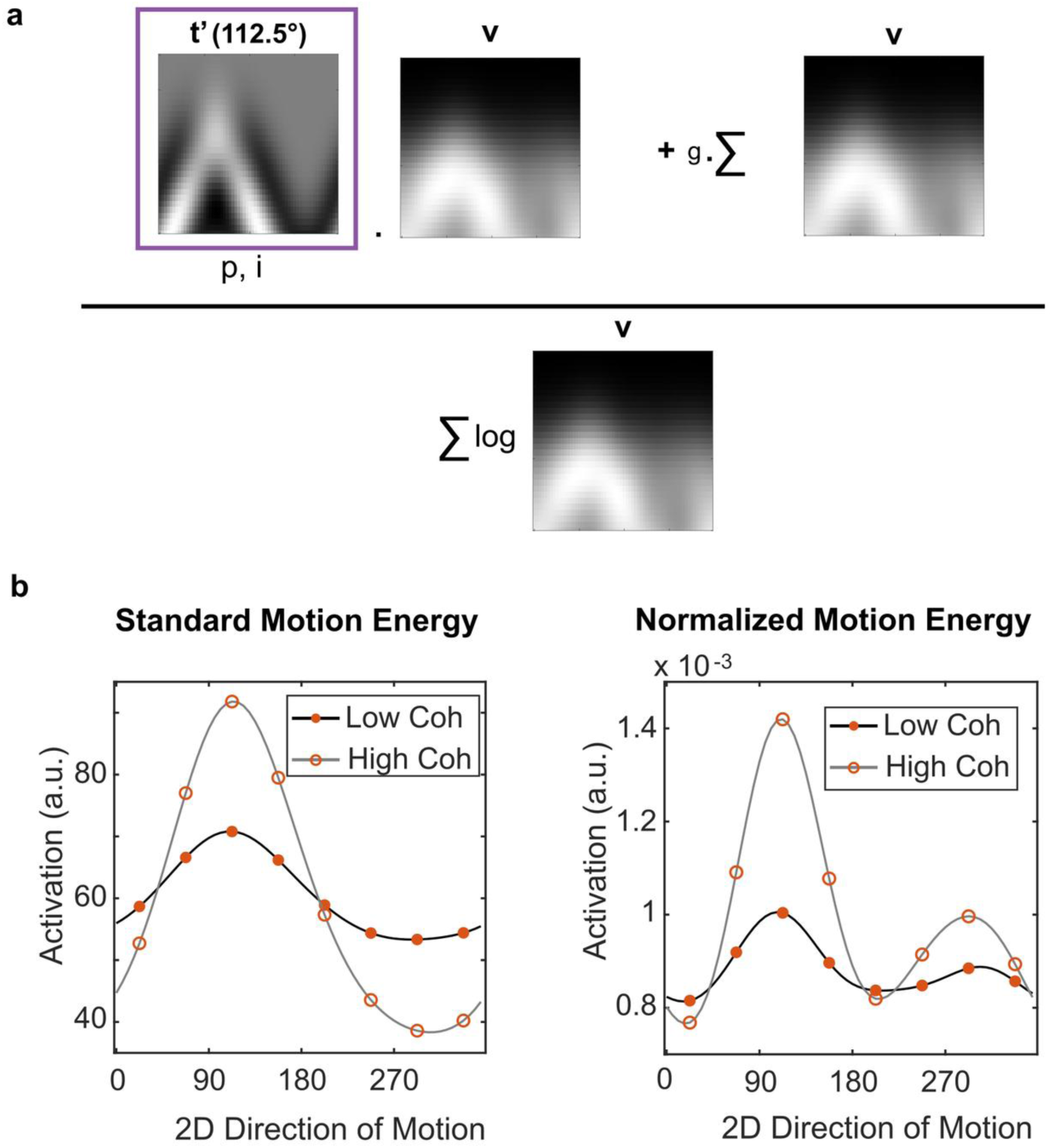
The normalized motion-energy (NME) model. (a) Two mechanisms are added to the standard model. First, each template is inhibited by the templates tuned to nearby directions, resulting here in the inhibited template for 112.5°; the inhibition is controlled by its total strength relative to excitation (p) and its tuning width (i). Second, an untuned gain control: a term proportional to the total motion energy, scaled by g, is added to the dot product between the inhibited template and the motion energy of the stimulus, and the result is divided by the summed log-compressed motion energy. Because this denominator is larger when motion energy is spread out, as at low coherence, overall activation increases with coherence. (b) Activation profiles of the two models when the true direction is 112.5°, at low and high coherence. In the NME model, tuned inhibition produces a secondary peak in the opposite direction, and total activation increases with coherence, whereas it is the same at both coherence levels in the standard model.

We next asked whether confidence judgments can be predicted directly from the perceptual signal, and which perceptual computations are required. In both models, confidence is read out from the same signal as the perceptual decision. Models were fitted in two stages, first to perceptual decisions and then to confidence judgments (see Evaluating models).

#### Perceptual decisions

We first fitted both models to the response proportions of the average observer (8 response categories relative to the true direction × 4 coherence levels). The SME model achieved an RMSE of 0.0491 (68% CI [0.0432, 0.0558]), whereas the NME model achieved a much lower RMSE of 0.00879 (68% CI [0.00862, 0.0120]; Fig 8). Qualitatively, the SME model failed in three ways: its accuracy increased too shallowly with coherence, it overpredicted errors to the adjacent directions (0.209 vs. 0.134 per alternative at 30% coherence), and it predicted almost no opposite-direction errors (0.00793 vs. 0.132). More generally, the SME model predicts errors that decrease monotonically with the distance from the true direction, whereas the data show a minimum at the orthogonal directions and a rise toward the opposite direction.

**Fig 8.**
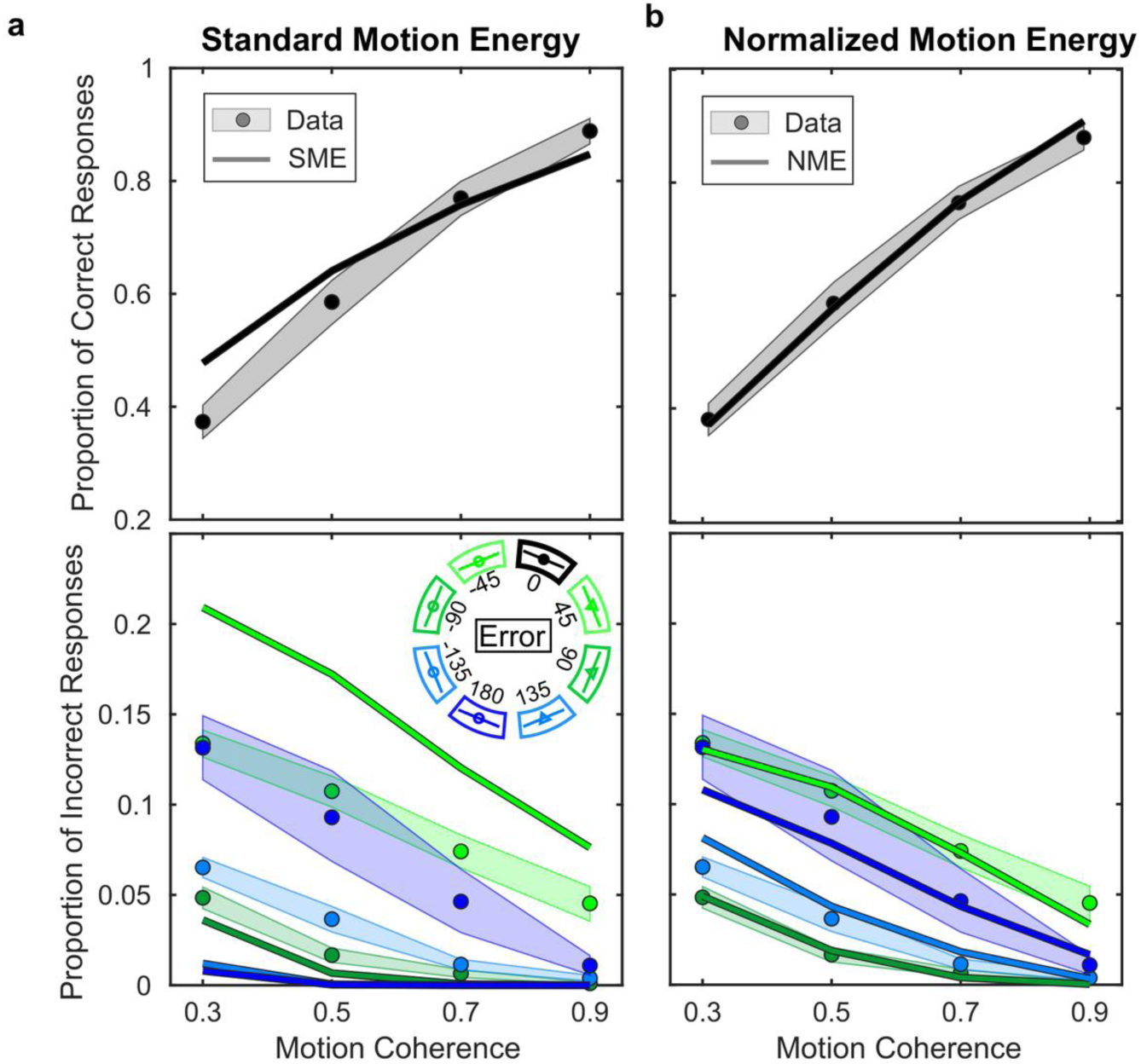
Models fit to perceptual decisions. (a) Standard motion-energy (SME) model; (b) normalized motion-energy (NME) model. Top: proportion of correct responses as a function of motion coherence. Bottom: proportion of errors per response alternative as a function of motion coherence, for errors of ±45° (light green), ±90° (dark green), ±135° (light blue) and 180° (dark blue); the two alternatives at the same angular distance (e.g., +45° and −45°) are averaged. Circles show the data of the average observer and shaded bands their 68% bootstrapped confidence intervals; lines show model predictions at the best-fitting parameters.

All fitted parameters fell within plausible ranges (Table 1). The decision noise corresponded to 0.203 (68% CI [0.184, 0.221]) of the range between minimum and maximum activation for the SME model, and to 0.353 (68% CI [0.274, 0.508]) for the NME model, averaged across trials and coherence levels. Inhibition strength, tuning width and decision noise trade off strongly, so the data alone constrain them only weakly: stronger and less sharply tuned inhibition improved the perceptual fit only slightly and left the confidence fit almost unchanged (see Evaluating models). We therefore fixed the inhibition strength of the NME model at 1.01, close to but slightly above excitation (set to 1), as the biologically most plausible solution, because suppressive inputs to MT neurons are comparable in strength to excitation [25]; this choice does not affect our conclusions. A strength of exactly 1 fitted worse (RMSE 0.019, adjusted R^2^ 0.992), so inhibition must slightly exceed excitation. With the strength fixed, the tuning width of inhibition was 10.0° (68% CI [9.0°, 11.8°]; 95% CI [8.3°, 14.3°]), so inhibition is sharply tuned if its strength is close to that of excitation. Whatever its strength, inhibition had to be tuned: with untuned inhibition, that is equal in all directions, the model failed as badly as the SME model (RMSE 0.045). The data therefore show that inhibition is centered on each channel’s preferred direction and weakest at the opposite direction.

**Table 1.** Best-fitting parameters of the SME and NME models for the average observer.

| Processing Stage | Parameter | SME Fit [68% CI] | NME Fit [68% CI] |
|---|---|---|---|
| **Perceptual** | Decision Noise (*σ*) | 0.203 [0.184, 0.221] | 0.353 [0.274, 0.508] |
|  | Inhibition Strength (*p*) | -- | 1.01 (fixed) |
|  | Inhibition Tuning Width (*i*) | -- | 10.0° [9.0°, 11.8°] |
| **Confidence** | Decay Rate (*k*) | 0.134 [0.051, 0.169] | 0.140 [0.020, 0.175] |
|  | Interval Bias (*b*) | 1.00 [0.993, 1.02] | 1.05 [0.987, 1.10] |
|  | Untuned Gain (*g*) | -- | 0.404 [0.227, 0.870] |

Perceptual stage: σ, decision noise added to the activation of each direction template, expressed as a fraction of the range between minimum and maximum activation (averaged across trials and coherence levels); p, total strength of inhibition relative to excitation, fixed at 1.01; i, tuning width (standard deviation) of inhibition. Confidence stage: k, rate at which the first-interval signal decays toward the average strength; b, multiplicative bias on the second-interval signal (b = 1: no bias; b > 1: bias toward the second interval); g, untuned gain, expressed as the mean value of the gain term (the second term of the numerator in Equation 6, divided by the denominator) as a proportion of the range between the minimum and maximum activation of the eight response channels across all simulated trials. Brackets: 68% confidence intervals from 2,000 bootstrap resamples of participants. Values are for the models without confidence noise.

Both models explained most of the variance in the data. Despite its additional parameter, the NME model had a higher adjusted R^2^ than the SME model (0.998 vs. 0.946) and a much lower AICc (−299 vs. −191; ΔAICc = 108, 95% CI [78, 115]). The NME model was preferred in all 2,000 bootstrapped datasets. It also provided a better fit than the SME model for every individual participant (S1 Text and S1 Table). A model based on linear vector averaging of the local motion signals, instead of IOC templates, did not change this conclusion: like the SME model, it failed to produce opposite-direction errors (S6 Fig). Thus, tuned inhibition is necessary to explain human global-motion discrimination performance.

#### Confidence judgments

We then fitted both models to the confidence judgments of the average observer, minimizing the RMSE between predicted and observed frequencies of first- and second-interval choices across all combinations of motion coherence and perceptual accuracy (4 × 4 × 4 × 2 categories; S8 Fig). The SME model achieved an RMSE of 0.00254 (68% CI [0.00254, 0.00294]) and the NME model a lower RMSE of 0.00183 (68% CI [0.00180, 0.00224]; Fig 9). Qualitatively, the SME model predicted a much sharper decline in first-interval choices as the perceptual error in the first interval increased, and it did not capture the relatively high confidence associated with opposite-direction errors (see also S2 Fig, where the data and both models are shown as choice matrices). Across coherence levels, the SME model also predicted a wider range of choice proportions and steeper slopes than observed. Both models reproduced the decline of first-interval choices along the equal-coherence diagonal (Fig 9, bottom).

**Fig 9.**
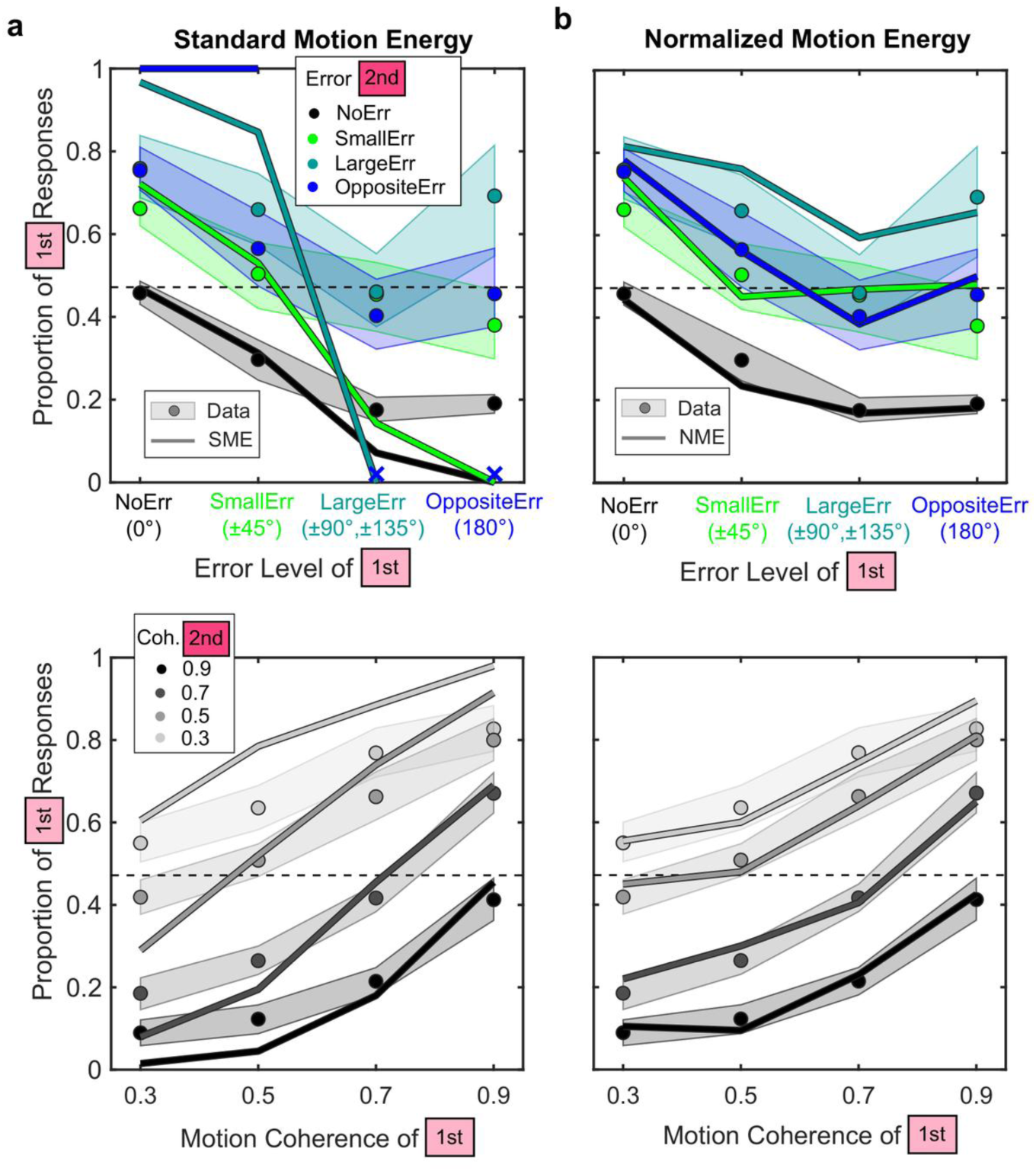
Model fits to confidence judgments. (a) Standard motion-energy (SME) model; (b) normalized motion-energy (NME) model. Top: proportion of first-interval confidence judgments as a function of the perceptual error in the first interval; colors indicate the error in the second interval (no error, black; small, green; large, cyan; opposite, blue). Bottom: proportion of first-interval choices as a function of motion coherence in the first interval; shades of gray indicate coherence in the second interval, from light (0.3) to black (0.9). Circles show the data of the average observer and shaded bands their 68% bootstrapped confidence intervals; lines show model predictions at the best-fitting parameters. The dashed line marks the overall proportion of first-interval choices. Crosses mark categories in which the model produced no trials (large and opposite-direction errors in the SME model). Predictions are for the models without confidence noise.

The best-fitting parameters fell within plausible ranges (Table 1). Both models showed a partial decay of the first-interval signal toward the overall mean (k = 0.134 for the SME model and 0.140 for the NME model), which accounts for the order bias observed when the two intervals had the same coherence. At the group level, neither model showed a meaningful interval bias (b = 1.00 and 1.05, where b = 1 indicates no bias).

Both models explained most of the variance in the confidence judgments. Despite its additional parameter (the untuned gain g), the NME model had a higher adjusted R^2^ than the SME model (0.944 vs. 0.893) and a lower AICc (−1,607 vs. −1,526; ΔAICc = 81.1, 95% CI [34, 119]), and it was preferred in 99.95% of bootstrapped datasets. These results were consistent across individuals: the NME model fitted clearly better for eleven of the twelve observers, and the SME model for one (S1 Text). As an additional analysis, we tested whether noise at the confidence stage contributes (see Evaluating models). We found that such noise is probably present and is about as large as the decision noise: its best-fitting standard deviation was 1.0 times the decision noise σ (68% CI [0.33, 1.03]). It nevertheless added little: the fit of the NME model improved only slightly (RMSE 0.00142 to 0.00131, adjusted R^2^ 0.966 to 0.971, ΔAICc = 17.4), and the untuned gain remained necessary (ΔAICc ≈ 65 when removed). Its main contribution concerned a quantity that was not part of the fit: without noise, the NME model overestimated the accuracy difference between selected and unselected intervals at a given coherence (12.8 vs. 7.8 percentage points in the data; Fig 5a), whereas with noise it predicted 8.2 percentage points. The NME model reproduced the direction of the preference for opposite over large errors in direct comparisons (Fig 9b). Together, these results show that tuned inhibition accounts for the direction of this confidence effect, and that the untuned gain, which lets total activation increase with motion coherence, captures the relationship between confidence and coherence better than the SME model, in which total activation is constant across coherence levels. Noise at the confidence stage is probably also present, at a size comparable to the decision noise, but it adds little to the fit beyond these two mechanisms.

Overall, two biologically plausible mechanisms accounted for both perceptual decisions and confidence judgments: tuned inhibition among direction channels produced the frequent opposite-direction errors and the direction of their confidence advantage, and an untuned gain control, which lets total activation increase with motion coherence, was needed for confidence to depend on coherence as strongly as observed.

## Discussion

To understand how confidence judgments are formed in a perceptual task, we used a multiple-aperture motion stimulus in which reporting the global motion direction requires spatially integrating locally ambiguous signals. Direction discrimination depended on motion coherence, as previously reported [14,18]. At the perceptual level, participants frequently chose the direction opposite to the true motion, and, in direct comparisons, assigned these errors more confidence than other large errors. At the metacognitive level, confidence tracked perceptual reliability beyond what the nominal coherence of the stimulus could explain, ruling out a simple stimulus-based heuristic. To identify the computations underlying these observations, we compared two image-computable models of motion integration. A normalized motion-energy model, incorporating tuned inhibition and an untuned gain control, accounted for both perceptual decisions and confidence judgments. In this model, confidence is read out directly from the perceptual signal, without a separate heuristic. Noise at the confidence stage, a component of current models of confidence [28,29], was probably also present: it added little to the overall fit, but it accounted for the accuracy difference between selected and unselected intervals. This suggests that confidence in this task relies on the perceptual signal.

### Perceptual mechanisms of opposite-direction errors

At the perceptual level, our findings contribute to a recent line of work on visual motion processing. Studies using continuous direction reports have revealed robust opposite-direction errors for random-dot kinematograms [30–34]. Similar errors may be present in multiple-aperture stimuli (see the error distributions in Fig 9 of [18]), but they have not been quantified. Here, we quantified them with an eight-alternative forced-choice task and found that, in direct comparisons, participants assigned more confidence to opposite-direction errors than to other large errors, although less than to small errors. This suggests that these errors are not simply guesses or motor errors, the latter being unlikely in any case because the opposite response arc lies on the far side of the response ring. Given their prevalence, some opposite-direction errors may have gone unreported in previous paradigms [35].

The neural mechanisms underlying these errors remain debated, but our model shows that strong, tuned inhibition among direction channels can account for them, and that this inhibition must be sharply tuned if its strength is close to that of excitation. Because the inhibition is centered on each channel’s preferred direction, it suppresses channels near the true direction most strongly and spares the opposite one, which then forms a secondary peak in the population response (Fig 7b). As coherence decreases, the difference between the two peaks shrinks, so that decision noise more often favors the opposite direction. The model thereby reproduces not only the frequency of opposite-direction errors but also their steeper decline with coherence compared with other large errors. Biologically, suppression between direction-selective units is a well-documented feature of V1 and MT [23,36], and the total strength of suppressive inputs to MT neurons is comparable to that of excitation [25], consistent with the near balance of excitation and inhibition in our reported model. How sharply this suppression is tuned in direction, however, remains open: Inagaki et al. [25] found it to be broadly distributed in the spatiotemporal frequency domain, whereas our model requires it to be concentrated around each channel’s preferred direction. Moreover, some random-dot studies did not report a dependence of opposite-direction errors on coherence [31,33]. Further work is needed to determine whether this discrepancy reflects the response procedure (discrete eight-alternative choices rather than continuous reports) or the local motion signals (1D oriented Gabors rather than 2D moving dots).

Other low-level signals may also contribute to the bimodal response. Spatiotemporal filtering produces some motion energy in the direction opposite to each Gabor’s drift (about 22% of the peak energy in our filters; S5 Fig, panel b), which our Gaussian approximation omits, and temporal smearing of this kind has been proposed for random-dot kinematograms [31,33]. In contrast, motion streaks cannot explain our results. Recent fMRI work proposed that bimodal direction representations arise from the ambiguous streaks left by fast-moving dots ([34]; see also [37–39]). In our stimuli, the Gaussian envelopes of the Gabors were stationary, so that no element changed position and no streaks could form; moreover, their drift speed (0.5 dva/s) was far below the critical speed of about 2.5 dva/s required for streaks to contribute to motion perception [37,39,40]. It will be important to assess the relative contributions of tuned inhibition, residual motion energy and motion streaks with faster-moving stimuli.

### Confidence formation in global motion direction decisions

Beyond the origin of the perceptual errors, our behavioral data provide insights into how confidence itself is formed. Our two-interval design introduced a known contraction bias, well explained by Bayesian models in which a decaying memory of the first stimulus is combined with a prior centered on the average of the experiment [41,42]. Our models accounted for this bias with a single decay parameter. Apart from it, confidence judgments closely tracked both the magnitude of perceptual errors and motion coherence. Unlike in random-dot kinematograms, the local elements of our stimuli require spatial integration to recover the global direction, so that no simple stimulus statistic is available as a proxy for confidence. Moreover, our data showed that observers did not rely on the nominal coherence of the stimulus alone, suggesting that perceptual reliability drives these judgments.

Our computational modeling supports this conclusion. In the NME model, tuned inhibition accounts for opposite-direction errors, and an untuned gain explains how confidence scales with motion coherence. Because the gain term is identical for all direction channels, it cancels out when the perceptual decision is made and affects only the comparison of confidence across stimuli of different coherence. The gain term is not a source of noise, however. The parameter g is fixed, and the quantity it multiplies, the total motion energy divided by the summed log-compressed energy, depends almost entirely on coherence: within a coherence level, the gain term varies across stimuli by only a few percent of the decision noise. The gain term therefore provides a nearly deterministic signal of stimulus coherence, and increasing g makes confidence judgments more strongly driven by coherence rather than closer to chance (S7 Fig). If observers were asked to judge coherence directly, this overall level of activation could serve as the primary perceptual signal.

Because this signal does not affect perceptual decisions, our behavioral data alone cannot determine whether it is part of the perceptual representation or computed separately for confidence. Several observations favor a perceptual origin. The mechanism resembles a compressive nonlinearity applied to the outputs of direction-selective units, whose sum is then used to normalize each pattern-motion unit; normalization by pooled activity is a canonical operation in motion and visual processing [21,43], although canonical models typically pool expansive rather than compressive responses. Moreover, the resulting increase of total activation with motion coherence matches the increase of total MT activity reported for random-dot kinematograms [26] and multiple-aperture stimuli [27]. A signal of this kind also agrees with recordings in V1, where a linear readout of population activity sufficed to predict perceptual decisions, whereas confidence was better predicted by a nonlinear readout of the same activity, whose output increased with overall response gain regardless of stimulus orientation [44]. Neural recordings during confidence judgments could test this directly: our model predicts that the coherence dependence of confidence should already be present in the overall activity of direction-selective populations.

More generally, the exact implementation of these mechanisms is less important than the phenomena they capture. Opposite-direction errors and the increase of total cortical activation with motion coherence are both established perceptual phenomena, and a model that reproduces them accounts for most of the variance in both our perceptual and confidence data. Our results therefore suggest that confidence in this task reflects the perceptual signal itself, including its complex structure, rather than a simplified summary of the stimulus. Observers may rely on heuristic strategies under other conditions, combining simple visual dimensions with higher-order cognitive factors [11,12,45], but in this global motion task confidence was clearly tied to perceptual reliability. This is consistent with neurophysiological recordings in primate lateral intraparietal cortex [46,47] and human EEG studies [48,49] showing a direct effect of perceptual reliability on confidence. In motion discrimination, microstimulation of MT alters confidence in a way that mimics a change in the visual motion [50], and both process models and recordings in lateral intraparietal cortex indicate that confidence draws on the same evidence as the perceptual decision [51,52].

### Confidence and Perceptual Reliability

Normative theories debate whether confidence represents a Bayesian posterior probability [7,8,53] or a measure of self-consistency [54]. Our mechanistic model does not distinguish between these accounts: confidence is based on the strength of the winning response after normalization, a quantity that could approximate either signal. Relying on the winning response agrees with recordings in the lateral intraparietal cortex, where the link between choice signals and confidence was found for neurons representing the chosen option [47,52], although behavior in the latter study was better explained by reading confidence from the losing option. We did not test whether confidence also depends on the gap to the second-best alternative, which best explained confidence in a three-alternative task [55], because reading out the winning response alone is simpler and already accounted for most of the variance in our data; this could be examined with eight-alternative tasks like ours. Human confidence also shows biases that appear suboptimal under these accounts, which may reflect computations optimized for different goals. For example, if observers implicitly evaluate a "no coherent motion" alternative, confidence would represent the probability of being correct in a higher-dimensional space rather than in a simple choice between alternatives [56].

Consistent with this view, recent work shows that seemingly suboptimal confidence behavior can emerge from basic properties of the underlying processing, such as tuned inhibition between decision units [57], the statistics of natural images [58] and the architecture of neural networks [59]. Our results extend this idea to the perceptual front end. Here, tuned inhibition operates between sensory direction channels in an image-computable model, rather than between decision units, and it accounts for a pattern that would otherwise look like a metacognitive failure: relatively high confidence in large, opposite-direction errors in direct comparisons. When such patterns can be explained by perceptual processing, invoking distinct, higher-level metacognitive mechanisms may be unnecessary. This underscores the value of image-computable models: by generating quantitative predictions from the stimulus itself, they can guide future neurophysiological and neuroimaging studies to determine how the brain computes confidence.

## Conclusion

In a global motion task that offered no simple stimulus statistic as a shortcut, confidence judgments tracked perceptual reliability, beyond what motion coherence alone could explain. An image-computable model of motion integration accounted for both perceptual decisions and confidence judgments with two perceptual mechanisms. Tuned inhibition among direction channels, sharp if it is close to excitation in strength, produced the opposite-direction errors. An untuned gain control increases overall activation with the coherence of the integrated motion signal. It leaves the perceptual decision unchanged, but at higher coherence it increases the strength of the percept, on which confidence is based. Confidence was read out directly from the same signal that drives the perceptual decision, and noise at the confidence stage, although probably present, added little beyond explaining the accuracy difference between selected and unselected intervals. Seemingly unusual metacognitive behavior, such as relatively high confidence in opposite-direction errors in direct comparisons and a strong dependence of confidence judgments on motion coherence, emerged from these perceptual computations rather than from a confidence-specific mechanism. Distinguishing true confidence mechanisms from byproducts of perception therefore requires modeling the perceptual front end explicitly, to determine what information it passes on to confidence.

## Materials and methods

### Experimental Methods

#### Stimulus design and display

All experiments and analyses were conducted using custom MATLAB code with the Psychophysics Toolbox [60,61]. Stimuli were presented on a gamma-corrected CRT monitor (Iiyama Vision Master Pro 454; 1280 × 1024 pixels, 60 Hz refresh rate) with 8-bit luminance resolution. At a viewing distance of 57.3 cm, the resolution was approximately 36.6 pixels per degree of visual angle (dva).

The global motion stimuli consisted of 468 drifting Gabor patches [14]. Each Gabor had a spatial frequency of 4 cycles/dva, a Michelson contrast of 40%, and a Gaussian envelope with a standard deviation of 0.14 dva (5 pixels), drawn within a 0.71 × 0.71 dva support. Gabors were assigned random orientations and phases, drawn independently for each presentation, and arranged on a square grid with a spacing of 0.63 dva. The grid filled an annular region with an outer diameter of 16 dva and a blank central disc of 4.4 dva in diameter.

On each presentation, the global motion direction was randomly selected from eight alternatives: 22.5°, 67.5°, 112.5°, 157.5°, 202.5°, 247.5°, 292.5° and 337.5°, where 0° denotes rightward motion and angles increase counterclockwise. These directions avoid the cardinal axes, along which direction discrimination is more precise [62,63], and place every alternative 22.5° from the nearest cardinal axis. The global speed was fixed at 0.5 dva/s. Motion coherence was set to one of four levels (30%, 50%, 70% or 90%) and determined the probability that each Gabor was a signal element. Signal elements drifted consistently with the global motion, whereas noise elements drifted consistently with global directions drawn independently from a uniform distribution between 0° and 360°, at the same global speed. Each element’s local 1D motion was then derived from its orientation: it drifted orthogonally to its orientation, at a speed equal to the global speed multiplied by the cosine of the angle between the global direction and the element’s normal.

A response ring surrounded the Gabor array, consisting of eight colored arcs corresponding to the eight possible global motion directions. Each arc spanned 35°, was centered on its direction and had a width of 0.4 dva. The ring was placed just outside the stimulus annulus, giving a total display diameter of approximately 18 dva.

#### Perceptual and confidence judgment task

Each trial lasted approximately 5 s and comprised two consecutive perceptual intervals followed by a confidence judgment (Fig 2a). In each interval, participants reported the perceived global motion direction of the stimulus; after both intervals, they indicated which of their two perceptual decisions was more likely to be correct. Each perceptual interval (Fig 2b) began with a central fixation dot presented for a random duration between 250 and 750 ms, immediately followed by the motion stimulus. The Gabors drifted at full contrast for 467 ms (28 frames), after which their contrast was linearly ramped down to zero over three frames (50 ms), during which they no longer drifted, to avoid an abrupt offset transient, giving a total stimulus duration of 517 ms (31 frames). At the stimulus offset, a mouse cursor appeared at the center of the screen, and participants reported the perceived global motion direction by clicking one of the surrounding arcs. When the cursor reached the ring, the arc at the cursor’s angle was highlighted (brighter, and widened to 0.6 dva) while the other arcs dimmed. After the click, the chosen arc brightened further for 200 ms to confirm the response.

The second interval followed the same structure, with independently sampled stimulus parameters (motion coherence, global direction, Gabor orientations and phases). All 16 combinations of motion coherence across the two intervals were presented equally often, and for each interval and coherence level, the eight global directions were presented equally often.

After the second perceptual decision, the fixation dot remained on the screen for 500 ms and then turned red, and text prompts for the first and second intervals appeared on the left and right of the screen, respectively (Fig 2b). Participants indicated which of their two decisions was more likely to be correct by pressing the left or right arrow key.

Each participant completed two experimental blocks, either in a single session lasting approximately two hours (six participants) or on separate days (six participants). The experiment began with an 18-trial training phase (the 9 coherence-pair combinations of 50%, 70% and 90% coherence, each repeated twice), during which participants received auditory feedback on both their perceptual decisions and their confidence judgments. Each block lasted about 45 minutes and was run without feedback. Each block was divided into eight sub-blocks separated by short breaks. In total, participants completed 40 repetitions of each of the 16 coherence-pair conditions, resulting in 640 experimental trials per participant.

#### Participants

Fifteen observers (six experienced and nine naive; ages 22–32; 10 female) completed the experiment. Three were excluded: one performed at chance level in motion direction discrimination, and two made confidence judgments at chance level. All analyses are based on the remaining twelve participants. All participants had normal or corrected-to-normal visual acuity and were compensated €10 per hour.

#### Ethics statement

The study was approved by the ethics committee Comité de protection des personnes Ouest V (Rennes, France) and conformed to the Declaration of Helsinki. Participants were recruited and tested between 27 July 2023 and 27 October 2023, and all provided written informed consent before taking part.

#### Data analysis

Behavioral results are reported for the average observer, computed from the trials pooled across the twelve participants (for Fig 5 and the statistics of accuracy and error rates, by averaging the proportions of the twelve participants), and the computational models were fitted to these pooled data. Confidence intervals were obtained by bootstrapping participants: we repeatedly resampled sets of twelve participants with replacement (10,000 resamples for the behavioral statistics and Fig 5; 2,000 for the error bars of the other figures and for model parameters). Figures, parameter estimates and fit statistics are reported with 68% confidence intervals, and differences between conditions or models with 95% confidence intervals.

We first checked that the data could be collapsed across intervals and motion directions. Perceptual performance did not differ systematically between the two intervals at the group level (65.5% vs. 65.4%; 5 of 12 participants performed better in the second interval, 6 in the first and 1 equally in both), so data were combined across intervals. We then checked for spatial biases. Participants tended to choose the two upper directions (67.5° and 112.5°) slightly more often than the two lower directions (247.5° and 292.5°), but this pattern was not consistent across individuals (6 of 12 observers). Conversely, normalized accuracy was higher for the lower than for the upper directions, again with limited consistency across observers. Because these biases were idiosyncratic, we collapsed the data across motion directions for all subsequent analyses.

### Modeling Methods

#### The standard motion-energy model (SME)

Computing motion energy directly from pixel-level input is straightforward for a single Gabor, but repeating it with a dense filter bank for all 468 elements of every stimulus is computationally prohibitive. We therefore implemented an efficient image-computable approximation, inspired by Kane et al. [24]: we first characterized the motion-energy signature of a single moving Gabor, and then used the relationship between this signature and the Gabor’s physical parameters to simulate the local motion energy of every element directly.

Motion-energy sensors were constructed following the Appendix of Kane et al. [24], with one modification: we swapped the cosine and sine functions defining the filter orientation, which shifts the zero-orientation reference. We used a single spatial-frequency channel matched to our stimuli (4 cycles/dva). Sensors and stimulus movies spanned 26 × 26 pixels (0.71 × 0.71 dva) and 30 frames (0.5 s). Unlike Kane et al. [24], we applied no temporal envelope; the sensors had only a spatial Gaussian envelope with a standard deviation of 5 pixels (0.14 dva), identical to that of the stimulus Gabors.

The filter bank comprised 64 directions spaced by about 5.7° and 31 temporal frequencies between 0 and 5 Hz, corresponding to phase shifts of 0° to 30° per frame. Because spatial frequency and filter size were fixed, these temporal frequencies map onto 1D speeds from 0% to 250% of the global stimulus speed (0.5 dva/s), allowing us to describe each filter by an effective preferred speed. Motion energy was computed as the quadrature sum of the responses of even- and odd-symmetric filters, each obtained by integrating the product of the stimulus and the filter over space and time.

To validate this pipeline, we computed the motion energy of a single drifting Gabor, as used in our task, and of two control stimuli: a plaid and a moving dot (S5 Fig). The motion energy of a single Gabor was well approximated by a 2D Gaussian in direction–speed space (R^2^ = 0.93), centered on the Gabor’s 1D speed and in the direction orthogonal to its orientation, as expected from the aperture problem. The standard deviations were 24° along the direction axis and 0.242 dva/s along the speed axis. The residual energy not captured by the Gaussian was not random: most of it lay in the opposite direction at slow speeds, reaching about 22% of the peak energy (S5 Fig, panel b). Our approximation therefore omits this weak opposite-direction energy, a point we return to in the Discussion.

These Gaussian parameters do not depend on the phase or contrast of the Gabor, but they do depend on its size and spatial frequency, which were identical for all elements in our stimuli. We could therefore use a single 2D Gaussian for all elements. The Gaussian wrapped around the circular direction axis, and energy at negative speeds was mapped onto the opposite direction (180° away) with the corresponding positive speed (Fig 6a). For each element, the Gaussian was evaluated within ±3 standard deviations of its mean and normalized to a maximum of one. Spatial integration was modeled as a linear sum of the local energies across all elements, giving the total motion-energy profile of the stimulus (Fig 6a).

For computational efficiency, we precomputed these total motion-energy profiles for 200 independent stimulus presentations in each of the 32 task conditions (8 global directions × 4 coherence levels), and used them for all model evaluations.

The multiple-aperture stimuli were specifically designed to probe the ‘aperture problem’ and its solutions [14]. In isolation, a single moving Gabor produces a perception of motion orthogonal to its orientation, corresponding to its peak in simulated motion energy. However, due to the aperture problem, this local 1D motion is consistent with multiple combinations of global (2D) directions and speeds. Crucially, for Gabors within the coherent set, these varying 1D motions are all mathematically consistent with a single rigid 2D pattern motion. Therefore, perceiving this global motion requires the visual system to integrate these ambiguous local signals across multiple apertures distributed in space.

While spatial motion integration can involve multiple processes [14,64–67], Amano et al. [14] showed that under conditions matching our experiment, integration follows a non-linear rule: the intersection of constraints provided by each Gabor (IOC [17,22]), rather than linear vector averaging [68–70]. Following Amano et al. [14], we therefore focused on IOC-based models. Because our task required only direction decisions, for which both rules predict the same mean direction, we also fitted a vector-average model in a separate analysis (S6 Fig).

Fig 6b illustrates this computation for a trial with a global motion direction of 22.5°, a speed of 0.5 dva/s and 30% coherence. For a 2D motion with direction *φ* and speed *V*, every signal element has a 1D velocity on the curve *s*(*θ*) = *V* ⋅ cos(*θ* − *φ*) in direction–speed space, where *θ* is the 1D direction of the element. This curve is the intersection-of-constraints (IOC) solution expressed in the space of the motion-energy filters: each point on it corresponds to one local constraint consistent with the global motion. The spatially integrated motion energy of the signal elements therefore concentrates along this half-cosine curve (Fig 6a, right), and recovering the global motion amounts to finding the 2D velocity whose curve best matches the integrated energy.

Although the IOC solution is mathematically well defined, the spread of motion energy around the curve depends on factors such as the sampling of local orientations and the tuning of the motion filters, which would be difficult to describe analytically. We therefore implemented IOC empirically, using templates in the spirit of Kane et al. [24]: the template for a given 2D direction is the average motion-energy profile of 100% coherent stimuli moving in that direction at the stimulus speed (Fig 6b, left). Each template thus represents the IOC curve of one 2D velocity, including the spread produced by the filters, and acts as a direction-selective unit tuned to that global motion, analogous to an MT pattern cell [21,23]. Because the global speed was fixed in our experiment, IOC was evaluated only at this speed, and the templates differ only in their preferred direction.

To build the templates, we averaged the motion-energy profiles of 700 presentations of 100% coherent stimuli moving at 22.5° and 0.5 dva/s. Templates for the other directions were obtained by circularly shifting this profile along the direction axis, giving 64 templates spaced by 5.625°, and each template was normalized to sum to one (Fig 6b). Perceptual decisions were based on the eight templates corresponding to the response alternatives; the full set of 64 templates is used for the tuned inhibition of the NME model (see below).

For each of the eight possible 2D directions *d*, the activation *r*(*d*) is the dot product of the template *t*(*d*) and the spatially integrated motion energy *v*, corrupted by independent Gaussian decision noise *ε* with zero mean and standard deviation *σ*, the only free parameter of the perceptual stage (Equation 1):

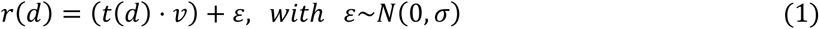

The perceived direction *d̂* is the direction with the maximum activation, and the strength of the percept *r̂* is that maximum activation (Equation 2):

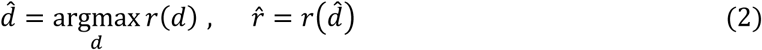

In each confidence trial, two perceived directions (*d̂*_1_, *d̂*_2_) are determined. We assume that confidence is directly related to the strength of the percept, so that the confidence judgment compares the strengths *r̂*_1_ and *r̂*_2_ of the two percepts. Two free parameters enter this comparison. First, because our behavioral data suggest that the first percept is partly forgotten, its strength decays toward the average strength across all trials and both intervals (*<u>r</u>*) at a rate *k*. Second, a multiplicative parameter *b* on the second interval accounts for interval biases (*b* = 1: no bias; *b* > 1: bias toward the second interval). The first interval is chosen (*c* = 1) when:

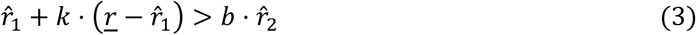

#### The normalized motion-energy model (NME)

The second model adds two mechanisms to the standard model. The first is inhibition among direction channels: each template is inhibited by the templates tuned to other directions. Suppressive inputs to MT neurons are strong, often comparable in total strength to excitation, and they sharpen velocity tuning [23,25]. Here, we assume that this inhibition is tuned around each channel’s preferred direction. This assumption differs from the motion-opponent suppression of Rust et al. [23] and from the broadly distributed suppression reported by Inagaki et al. [25]; it was chosen because it leaves the opposite direction largely uninhibited, and its tuning width is a free parameter estimated from the data. Inhibition acts across the full set of 64 templates, spaced by 5.625°, not only the eight response alternatives. The inhibited template for direction d is obtained by subtracting a weighted sum of all other templates:

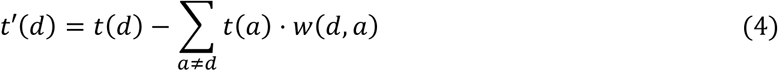

The weights follow a circular Gaussian of the angular distance *Δ*(*d*, *a*) between the two directions, excluding the template itself, and are normalized to sum to *p*:

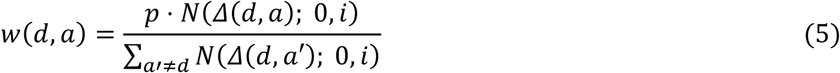

where *N*(·; 0, *i*) is a Gaussian probability density with zero mean and standard deviation *i*. The two new parameters thus control the total strength of inhibition relative to excitation (*p*, with excitation set to one) and its tuning width (*i*, expressed in degrees). With strong, tuned inhibition, as in the fits reported here, the inhibited templates contain large negative regions (Fig 7a), so that the dot product *t*′(*d*) ⋅ *v* is often negative.

The second mechanism, an untuned gain control, restores a positive baseline and makes overall activity depend on motion coherence. It builds on the untuned divisive normalization of motion-energy models of MT [21,23,43], consistent with the broadly distributed suppression observed in MT neurons [25]. It differs from these models in two ways, chosen to reproduce the increase of total MT activity with motion coherence [26,27]. First, an untuned excitatory term, proportional to the total motion energy of the stimulus (∑_*j*_ *v_j_*) and scaled by a free parameter *g*, is added to every direction channel. Second, the result is divided by the summed compressed rather than squared motion energy, ∑*_j_* log(*v_j_* + 1), where the sums run over all direction–speed bins of *v*; with an expansive pool, concentrated energy at high coherence would instead reduce total activity. The constant 1 is added to each bin before the logarithm, so that bins without energy contribute zero; log(*v_j_* + 1) thus acts as a compressive nonlinearity applied to each channel before pooling. Because this nonlinearity is compressive, energy spread over many bins (as at low coherence) yields a larger denominator than the same energy concentrated in fewer bins (as at high coherence), so that the overall response increases with coherence. Decision noise is then added to the normalized activations:

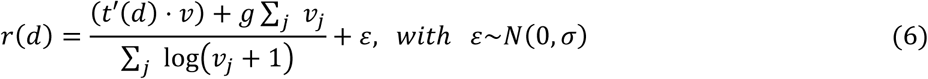

This sequence of computations is illustrated in Fig 7a. Once the activation is computed for each of the eight possible 2D motion directions, perceptual decisions and confidence judgments are made exactly as in the standard model, using Equations 2 and 3.

The impact of tuned inhibition depends on both parameters, p and i, which together shape the distribution of correct and incorrect responses. For example, when total inhibition slightly exceeds excitation and is sharply tuned, a secondary peak emerges in the activation profile at the opposite direction (Fig 7b), because the opposite-direction channel is far enough from the true direction to escape its narrow band of strong suppression.

The divisive normalization component (the sum of the logarithm) exerts a minor, fixed influence on perceptual decisions. Because this denominator scales the responses before the constant decision noise (*ε*) is added, it directly alters the effective signal-to-noise ratio. Specifically, at lower coherence levels, the widespread motion energy results in a larger denominator, which shrinks the differences in activation between alternative directions. Consequently, the fixed-variance decision noise has a greater relative impact, slightly increasing the probability that the maximum activation flips to an incorrect direction.

In contrast, the untuned gain, controlled by g, has no effect on perceptual decisions. The gain term, including its division by the summed log-compressed energy, takes the same value for every candidate direction within a stimulus, so it cancels out when the maximum response is selected; g therefore influences only confidence judgments. Increasing g does not act like noise at the confidence stage; instead, it makes confidence judgments sharper. With a larger g, confidence follows the coherence of the two intervals more strictly, pushing choices across coherence pairs toward all-or-none, whereas adding noise at the confidence stage pushes all choices toward chance (S7 Fig). This may seem surprising, because the gain term is the same for every direction and could therefore be expected to behave like confidence noise. However, it is not random. The parameter g itself is fixed; what changes from one stimulus to the next is the quantity it multiplies, the total motion energy divided by the summed log-compressed energy (Equation 6). This quantity is determined almost entirely by coherence: its mean increases with coherence, whereas across stimuli of the same coherence the resulting gain term varies by only 1–3% of the decision noise σ at the best-fitting parameters. The gain term therefore adds more to the strength of the percept in the higher-coherence interval, acting as a coherence signal rather than as random variability, and a larger g strengthens this signal.

#### Evaluating models

Both models were fitted to the data of each participant and of the average observer (data pooled across participants). Because perceptual decisions and confidence judgments depend on distinct parameters, fitting proceeds in two stages. Model predictions were obtained by simulation, using the 200 precomputed stimulus presentations of each of the 32 conditions (8 directions × 4 coherence levels). Confidence trials were formed by randomly pairing presentations within each of the 16 combinations of first- and second-interval coherence (200 pairs each). The random draws were fixed across evaluations, so that the fitting objective was deterministic. Parameters were optimized with the Nelder–Mead simplex method (*fminsearch*, MATLAB).

In the first stage, the models were fitted to perceptual decisions. The free parameters were the decision noise *σ* for the SME model, and *σ*, with the inhibition strength *p* fixed at 1.01, and the inhibition width *i* for the NME model. Because *g* does not affect perceptual decisions, it was fixed at this stage. The models were fitted to the proportions of each response relative to the true direction (correct, ±45°, ±90°, ±135° and 180°, with the two signs kept separate), at each of the four coherence levels, collapsed across the eight true directions (8 × 4 proportions). We minimized the root-mean-squared error (RMSE) between predicted and observed proportions, with predictions averaged over 25 independent random streams (40,000 simulated trials per coherence level). Because inhibition strength, tuning width and decision noise trade off strongly (see Modeling Results), the strength was fixed at 1.01, slightly above excitation: suppressive inputs to MT neurons are comparable in strength to excitation [25], and a strength of exactly 1 fitted poorly. For the NME model, predictions were computed on a grid (47 widths from 5° to 36° and 40 noise levels), and the best points were refined with the Nelder–Mead simplex method on the exact objective. For the SME and vector-average models, the decision noise was estimated on a fine grid.

In the second stage, the models were fitted to confidence judgments, with the perceptual parameters fixed at their best-fitting values. The free parameters were the decay rate *k* and the interval bias *b* for the SME model, and *k*, *b* and the untuned gain *g* for the NME model. Confidence judgments were sorted by the coherence of the two intervals (4 × 4) and by their joint perceptual accuracy (both correct, first correct and second incorrect, first incorrect and second correct, both incorrect), giving 4 × 4 × 4 categories. Because perceptual accuracy is a behavioral outcome rather than a controlled variable, trial counts were unevenly distributed across these categories. We therefore fitted the counts of both first- and second-interval choices in every category (4 × 4 × 4 × 2), each divided by the total number of trials in the corresponding dataset, so that model predictions and data were expressed as relative frequencies on the same scale. We again minimized the RMSE. Because the objective is piecewise constant, this stage was optimized from 28 starting points (8 for bootstrapped datasets), with the constraints g ≥ 0, 0 ≤ k ≤ 1 and b > 0, and the best solution was kept. Because the fit is nearly flat along k, each search was followed by a scan over k and a final search from the best point; the same was done for the noise in the models with confidence noise.

Models were compared with the adjusted R^2^ and with the corrected Akaike Information Criterion adapted for least-squares fitting, AICc = n·ln(RMSE^2^) + 2m + 2m(m + 1)/(n − m − 1), where n is the number of data points (32 for perceptual decisions, 128 for confidence judgments) and m the number of free parameters [71,72]. Confidence intervals for parameters and fit statistics were obtained by repeating the fitting procedure on 2,000 bootstrapped datasets, in which participants were resampled with replacement.

To test whether confidence requires variability beyond the perceptual signal, we also fitted a version of the NME model in which independent Gaussian noise, with a free standard deviation, was added to the strength of each percept before the comparison of Equation 3. Because the effect of this additional parameter can be similar in size to the sampling variability of the simulated trials, we repeated this comparison with 25 times more simulated trials to reduce simulation noise, pooling 25 independent random streams of noise and trial pairings (5,000 pairs per coherence combination), and evaluated the resulting fits on 25 further streams not used for fitting. To examine how inhibition strength, tuning width and decision noise trade off, we fitted the perceptual stage with the strength free. We computed predictions on a grid of 26 widths from 7° to 110°, with 11 strengths and 13 noise levels at each width, centered on the values that fitted best at that width, because these change strongly with the width. For the average observer, each participant and each of the 2,000 bootstrapped datasets, we selected the best grid point. The best fit of the average observer had a width of 50° and a strength of 1.34, a small improvement in absolute terms over the fixed-strength model (adjusted R^2^ 0.999 vs. 0.998; RMSE 0.00685 vs. 0.00879; ΔAICc ≈ 14). The best width had a 68% CI of [26°, 90°] and a 95% CI of [9°, 110°]. Untuned inhibition was approximated with very large widths (1,000° and above), with strength and decision noise optimized with the Nelder–Mead simplex method. Fits at other fixed strengths (1 to 1.05) followed the procedure described above: strengths from 1.005 to 1.03 gave widths from 6.6° to 17.7° with an adjusted R^2^ of 0.998. To test how the confidence fit depends on these parameters, we also fitted the confidence stage across widths, with strength and decision noise at their best values for each width, on 25 random streams, and evaluated it on 25 further streams. It was almost unchanged (RMSE on simulated trials not used for fitting: 0.00148 at 10° and 0.00143 at 50°). To test the IOC assumption, we fitted a vector-average model [14,68–70], in which the global motion is the linear average of the local motion signals rather than their intersection of constraints. Each direction–speed bin of the total motion-energy profile of the stimulus (Fig 6a) represents one velocity vector, and the model computes the average of these vectors, weighted by their motion energy. Independent Gaussian noise, whose standard deviation was the only free parameter, was added to the horizontal and vertical components of this average velocity, and the perceived direction was the response alternative closest to the direction of the resulting vector. The noise was estimated with the same procedure as for the perceptual stage of the other models.

## Data availability

All data and analysis code (MATLAB) underlying the findings of this study are available on Zenodo at https://doi.org/10.5281/zenodo.23993199 (data: CC BY 4.0; code: MIT License). The files will be made public upon publication.

## Supporting information

### S1 Text. Individual participant model fits

**S1 Table.** NME parameters and model comparison (ΔAICc = NME − SME; negative favors NME). i, inhibition width, with strength fixed at 1.01; k, decay rate; b, interval bias; g, untuned gain in raw model units (Table 1 gives g for the average observer as a proportion of the activation range). Participants are numbered as in S1–S4 Figs.

**S1 Fig.** Perceptual errors of individual participants. Proportion of incorrect responses per response alternative as a function of motion coherence, for each participant (format of Fig 3b): ±45° (light green), ±90° (dark green), ±135° (light blue) and 180° (dark blue); the two alternatives at the same angular distance are plotted separately (circles and triangles).

**S2 Fig.** Confidence judgments as matrices, for the data and both models. Each cell shows the proportion of trials in which the first interval was chosen as more likely to be correct (gray scale from 0, black, to 1, white). Rows correspond to the first interval and columns to the second. Top row: proportions by perceptual error category in each interval (no error, 0°; small, ±45°; large, ±90° and ±135°; opposite, 180°). Columns are ordered from opposite error to no error, so that the highest proportions appear in the top-left corner. Bottom row: proportions by motion coherence in each interval (0.3 to 0.9). Left: human data (average observer); same data as in **Fig 4**. Middle and right: predictions of the standard (SME) and normalized (NME) motion-energy models at their best-fitting parameters; same predictions as in **Fig 9**. The SME model predicts nearly all-or-none choices across error combinations, whereas the NME model reproduces the graded pattern of the data, including the relatively high proportion of choices for intervals with opposite-direction errors. Crosses mark categories in which the model produced no trials.

**S3 Fig.** Confidence judgments of individual participants as a function of perceptual error. Proportion of first-interval choices as a function of the error in the first interval, for each error category in the second interval (format of **Fig 4a**): no error (black), small (green), large (cyan) and opposite (blue). Marker size is proportional to the number of trials; missing points are combinations without trials. Dashed line: overall proportion of first-interval choices of that participant.

**S4 Fig.** Confidence judgments of individual participants as a function of motion coherence. Proportion of first-interval choices as a function of coherence in the first interval, for each coherence in the second interval, from light gray (0.3) to black (0.9) (format of Fig 4b).

**S5 Fig.** Motion-energy signatures of single stimuli. Motion energy as a function of the preferred direction and speed of the filters (64 directions, 31 speeds from 0 to 1.25 dva/s, 4 cycles/dva, Gaussian envelope SD 0.14 dva, 26 × 26 pixels × 30 frames). Insets: one frame of each stimulus; the red arrow gives its motion direction. (a) Control stimuli. Left: a plaid made of two gratings whose orientations differ by 120° (component speed 0.825 dva/s, pattern speed 1.65 dva/s); the energy forms two peaks, one per component, as expected for 1D motion sensors. Right: a moving dot (Gaussian blob, SD 2.5 pixels) at 0.825 dva/s; the energy forms an arch across directions, peaking at the dot’s direction (cf. [24]). (b) A single drifting Gabor as used in the experiment (0.5 dva/s, direction 180°). Left: computed energy. Right: best-fitting 2D Gaussian (SD 24° in direction and 0.242 dva/s in speed; R^2^ = 0.93), used to simulate every element. The residual energy lies mainly in the opposite direction (here 0°) at slow speeds, reaching about 22% of the peak.

**S6 Fig.** Vector-average model compared with the IOC-based models. Fits to the perceptual decisions of the average observer for a model in which the perceived direction is the alternative closest to the energy-weighted vector average of the local velocities (left), and for the SME (middle) and NME (right) models. Top: proportion correct. Bottom: errors per response alternative (colors as in **Fig 3b**). Solid lines: data with 68% bootstrapped confidence intervals; dashed lines: model predictions. The vector-average model fitted slightly better than the SME model (ΔAICc = −7.9, 95% CI [−11.1, −4.9]) but much worse than the NME model (ΔAICc = 100, 95% CI [71, 107]), and like the SME model it predicted almost no opposite-direction errors.

**S7 Fig.** The untuned gain is not confidence noise. Confidence judgments predicted by the NME model with all parameters at their best fit to the average observer, except, in the top two rows, the untuned gain g, set to 0, 1, 3 and 10 times its best-fitting value, and, in the bottom two rows, Gaussian noise added to each percept strength at the confidence stage, with SD 0, 0.5, 1 and 2 times the decision noise σ. Rows 1 and 3: first-interval choices as a function of the error in the first interval (format of **Fig 4a**); rows 2 and 4: as a function of coherence in the first interval (format of **Fig 4b**). Increasing g spreads choices across coherence pairs (range from 0.80 at the best fit to 1.00 at 10 times the best fit), whereas confidence noise compresses them toward chance (range 0.29 with noise of 2σ). At the best fit, the gain term varies by only 1–3% of σ across stimuli of the same coherence.

**S8 Fig. Confidence judgments by joint perceptual accuracy. Proportion of first-interval choices of the average observer as a function of coherence in the first interval, for each coherence in the second interval (shades of gray as in Fig 4b), separately for trials in which both decisions were correct (C,C), only the first (C,I), only the second (I,C), or neither (I,I). The confidence stage of the models was fitted to the choice frequencies in these 4 × 4 × 4 × 2 categories. Marker size is proportional to the number of trials; error bars show 68% bootstrapped confidence intervals; dashed line: overall proportion of first-interval choices (0.47)**.

